# Why Blood Pressure and Body Mass Should be Controlled for in Resting-State Functional Magnetic Resonance Imaging Studies

**DOI:** 10.1101/2021.06.02.446721

**Authors:** Guro Stensby Sjuls, Karsten Specht

## Abstract

Replicability has become an increasing focus within the scientific communities with the ongoing “replication crisis”. One area that appears to struggle with unreliable results is resting-state functional magnetic resonance imaging (rs-fMRI). Therefore, the current study aimed to improve the knowledge of endogenous factors that contribute to inter-individual variability. Arterial blood pressure, body mass, hematocrit, and glycated hemoglobin were investigated as potential sources of between-subject variability in rs-fMRI, in healthy individuals. Whether changes in resting state-networks (rs-networks) could be attributed to variability in the BOLD-signal, changes in neuronal activity, or both, was of special interest. Within-subject parameters were estimated utilizing Dynamic Causal Modelling (DCM) as it allows to make inferences on the estimated hemodynamic (BOLD-signal dynamics) and neuronal parameters (effective connectivity) separately. The results of the analyses imply that blood pressure and body mass can cause between-subject and between-group variability in the BOLD-signal and that all the included factors can affect the underlying connectivity. Given the results of the current and previous studies, rs-fMRI results appear to be susceptible to a range of factors, which is likely to contribute to the low degree of replicability of these studies. Interestingly, the highest degree of variability seems to appear within the much-studied Default Mode Network and its connections to other networks.

## Introduction

When studying the resting brain by utilizing resting-state functional magnetic resonance imaging (rs-fMRI), the brain’s energy consumption is measured as the participant is resting in a MR-scanner (Biswal, Zerrin Yetkin, Haughton & Hyde, 1995). This activity fluctuates in a range below 0.1Hz, and is hypothesized to support communication among neurons in the absence of a specific task (Biswal et al., 1995; Raichle & Mintun, 2006). The measured time-series of the fluctuations can further be analyzed and organized into fairly consistent resting-state networks (rs-networks) (Allen et al., 2011).

### Resting-State Networks

The growth of rs-fMRI studies over the last decades can in part be explained by the discovery of a consistent, functionally connected pattern of distributed brain regions; as a persistent network of “deactivation”. The activation of this Default Mode Network (DMN) ceases when a goal-directed or attention-demanding behavior or task is initiated (Raichle et al., 2001). While there is no unified view of the function of DMN as of yet, it has been postulated to support a “default mode” of the brain when an individual is awake and alert, but not actively involved in a task (Raichle et al., 2001). Others have suggested DMN to be involved in a self-referential and introspective state (Greicius, Krasnow, Reiss & Menon, 2003; Singh & Fawcett, 2008), involved in mediating the processes where one, for example, retrieves memories, plan for the future, or processing of one’s own impressions and feelings (Buckner, Andrews-Hanna & Schacter, 2008).

As DMN is “deactivated” when a cognitive task is performed, the activation of the Central Executive Network (CEN) is increasing, and the anti-correlation between DMN and CEN have been shown to increase with the degree of task difficulty (Fox et al., 2005; Hugdahl, Raichle, Mitra & Specht, 2015; Hugdahl et al. 2019). CEN is a task-related network, and it is believed to be involved in the manipulation and maintenance of information in working memory (Bressler & Menon, 2010; Toro, Fox & Paus, 2008). It is further postulated to be involved in decision-making in goal-directed behavior, attention, response inhibition and other executive functions, which qualitatively separates it from DMN (Bressler & Menon, 2010; Koechlin & Summerfield, 2007).

The Salience Network (SN) is involved in bottom-up detection of salience, by directing attention and memory resources to salient events. It has been shown to play a mediating role in up- and downregulation of DMN and CEN (Sridharan, Levitin & Menon, 2008), as a possible link between stimulus-driven processing and monitoring the internal environment of the brain and body (Craig, 2009; Menon & Uddin, 2010).

### Endogenous Sources of Variability in rs-fMRI

Despite its recent rise in popularity, the results of rs-fMRI seem to show between- and within-subject variability with a variety of endogenous and exogenous factors (Specht, 2019). Given that the BOLD-signal measured with fMRI is merely *related* to neuronal activity, and actually arises from a combination of changes in cerebral blood flow (CBF), cerebral blood volume (CBV), and oxidative metabolism to meet the energy demands of the active brain (Gauthier & Fan, 2019), other factors that can contribute to changes in CBF and CBV, e.g. arterial blood pressure, body mass and fat, and the velocity of the blood, might impact the BOLD-signal, and in turn the rs-fMRI results (Buxton, Wong & Frank, 1998). Several mathematical models of the hemodynamic response have been proposed to better understand and make predictions about the intricate relationship between neuronal and hemodynamic responses. Among these are the non-linear Balloon model (Buxton et al., 1998), which treats the venous compartments as a balloon that inflates due to increased CBF. The CBV therefore increases, and as a consequence deoxygenated hemoglobin (dHb) is released at a faster rate. In turn, this affects the BOLD-signal, essentially prolonging it (Buxton, Uludağ, Dubowitz & Liu, 2004). Factors that can affect CBV and CBF, like body mass (BMI) and blood pressure (BP), are put forward in the model as affecting the BOLD-signal (Buxton, 2012).

In addition to possibly affecting the BOLD-signal dynamics, some studies also imply that the connectivity between and within rs-networks can change with several endogenous factors. The relationship between BP and connectivity in the rs-networks, in normotensive individuals, has to the best of the authors knowledge not been investigated. However, several studies have been conducted on individuals with hypertension, employing both rs-fMRI, rat models and functional near-infrared spectroscopy (fNIRS) (Bu et al., 2018; Carnevale et al., 2020; Huang et al., 2016). Hematocrit-levels (HCT) has been associated with regional differences in functional connectivity. Regions associated with DMN, CEN and SN, namely anterior cingulate cortex (ACC), and medial prefrontal cortex (mPFC), intraparietal sulcus, insula, and opercular cortex, have been found to show between-subject variation with HCT levels (Yang, Craddock & Milham, 2015). Yang et al. (2015) rightly point out that it is unclear whether these differences are due to neuronal or non-neuronal variation. Some studies have found higher BMI to be related to decreased within-network connectivity of DMN, CEN and SN, as well as increased between-network connectivity (Chao et al., 2018; Doucet, Rasgon, McEwans, Micali & Frangou, 2017; Sadler, Shearrer & Burger, 2018). However, the regions included in these studies, as well as the results, vary. Still, several of the authors hypothesize that higher BMI is linked to changes in networks that balance sensory-driven and internally-guided (CEN, DMN) states; this might lead to weight gain as a consequence of poorly regulated eating behavior (Doucet et al., 2017; Sadler et al., 2018). Glycated hemoglobin (HbA1c) and changes in rs-fMRI connectivity have been studied, by comparing pre-diabetic or diabetic individuals with healthy individuals (Sadler, Shearrer & Burger, 2019). Similarly to the conclusions drawn from the previously mentioned BMI results, the authors discuss whether differences between groups are associated with differences in self-control; the functional connectivity pattern of the healthy individuals show stronger functional connectivity between a ventral attention network and a cingulo-opercular network, whereas the functional connectivity pattern of prediabetic individuals have been found to be stronger between a ventral attention network, a visual and a somatosensory network (Sadler, Shearrer & Burger, 2019). Individuals with type 2 diabetes mellitus have also been found to exhibit weaker functional connectivity in the right insula, and from the right insula to the bilateral superior parietal lobule, compared to healthy individuals (Liu et al., 2017).

### The Replication Crisis in the Field of Resting-State fMRI

As BP, HCT, BMI and HbA1c possibly affect cerebral hemodynamics, which could affect and disturb the BOLD-signal, it is uncertain whether the previous results on functional connectivity really represents changes in neuronal activity. If the results can be ascribed to changes in cerebral hemodynamics, the results mentioned above might have been wrongly attributed to changes in neuronal activity. In effect, rs-variability relates to the ongoing replication crisis in the field of psychological and medical research. Several studies indicate that rs-fMRI studies seem to produce highly unreliable results; showing within- and between-subject variability with a range of different endogenous and exogenous factors. These include the time of year and the time of day, circadian rhythm, sleep duration, prior events, mood, age and gender, to mention only a few (Agcaoglu, Miller, Mayer, Hugdahl & Calhoun, 2015; Choe et al., 2015; Curtis, Williams, Jones & Anderson, 2016; Goldstone et al., 2016; Harrison et al., 2008; Hjelmervik, Hausmann, Osnes, Westerhausen, & Specht; Hodkinson et al., 2014; Waites, Stanislavsky, Abbott & Jackson, 2005). Arguably, these findings give rise to skepticism about the rs-networks presumed stability (Specht, 2019).

### Towards Higher Replicability within the Field of rs-fMRI

To further increase the knowledge on between-subject variability in the rs-networks of healthy individuals, the effect of BP, HCT, BMI and HbA1c was investigated in this study. A research procedure that allows for functionally separating the BOLD-signal variation that can be attributed to hemodynamics, and the BOLD-signal variation that can be attributed to neuronal activity, was considered as highly relevant: Firstly, if some of the variation ascribed to variance in functional connectivity might in fact be attributable to hemodynamic variance, the conclusions drawn from functional connectivity studies might be flawed. Secondly, if the variables cause variability in both the hemodynamic and neuronal parameters of the BOLD-signal, it would imply that at least some of the variation in the hemodynamic parameters should be accounted for when drawing conclusions on connectivity. Thirdly, if all of the potentially observed variability can be ascribed to the neuronal parameters of the BOLD-signal, it could potentially confirm previous studies on rs-connectivity. And finally, a study that aims to ascribe the potential variability caused by some endogenous factors to either the hemodynamic response independently of neuronal activity, or the neuronal activity in rs-networks, has not previously been conducted.

In the current study it was aimed at investigating the between-subject variability that BP, HCT, BMI and HbA1c might cause in large scale rs-networks, as well as the between-group variability potentially caused by BMI, in a healthy population. The overarching implications of the current study relate to the ongoing replication crisis and what measures can be taken to ensure more reliable results in the fast-growing field of rs-fMRI; essentially facilitating more reliable results for future studies.

### Hypotheses

It was hypothesized that increased BP, BMI and HbA1c will weaken the BOLD-signal, whereas increased HCT was hypothesized to strengthen the BOLD-signal. Further, it was expected to detect that increased HCT will weaken the internal connectivity of DMN, CEN and SN, and, similarly, that increased BMI will weaken the internal connectivity of DMN and SN, but increase the between-network connectivity. Finally, it was hypothesized that increased HbA1c values will weaken the internal connectivity of CEN and SN, but increase the between-network connectivity.

To investigate these hypotheses, a hierarchical between-subject and between-group (based on BMI) design was chosen, based on the parameter from a cross-spectral density dynamic-causal modelling (csd-DCM) analysis. Generally speaking, DCM is generative model that infers on hidden neuronal states and activity given the measured fMRI signal (Friston, 2009). For resting-state fMRI, this framework has recently been extended to infer on functional connectivity from the frequency domain, by parametrizing the spectral characteristics of the neuronal fluctuations (Friston, Kahan, Biswal & Razi, 2014; Razi, Kahan, Rees & Friston, 2015). Such a csd-DCM is superior to other methods in this context, since it allows examining the individual parameters of the hemodynamic response, effective connectivity, cross-spectral density parameters α (amplitude) and β (density of the neural fluctuations), and Free Energy (model evidence), and their relationship to the endogenous factors BP, HCT, BMI and HbA1.

## Methods

A large sample of healthy young subjects were studied, to give indications of variability within a normal population, and to ensure the study’s statistical power. After ethical and practical considerations, previously collected data from the Human Connectome Project (HCP) was considered sufficient to answer the research question. Information on the subjects and the data collecting protocols are available online, allowing transparency and insight into potential advantages and disadvantages of the data used (HumanConnectomeProject, 2017).

### Subjects

The data used in the current study stems from the HCP S1200 release, which consists of data from healthy subjects from families with twins, born in the Missouri area (U.S.) and ranging from 22 to 35 years of age (Van Essen et al., 2013). A subsample (N=594) was semi-randomly chosen on the basis of an equal distribution of gender and age, as a part of a study on time-of-day effects on rs-fMRI effective connectivity and hemodynamic response. Therefore, the participants were also equally distributed throughout the day, in terms of scanning times (from 09.00 to 21.00) (for details, see Vaisvilaite, Hushagen, Grønli & Specht, 2020). Individuals with diabetes and high BP were excluded from the HCP data collection “as these might negatively impact neuroimaging data quality”. However, subjects with undiagnosed high BP and diabetes kept under control by means of diet were included in the study. The study sample (N = 594) was similar to the total HCP sample (N = 1206) in terms of BP, HCT, BMI and HbA1c, in addition to having a similar distribution of gender and age; full sample (N = 1206; female = 656, male = 550, mean age = 28,8; SD = 3.6), study sample (N = 594, female = 310, male = 284, mean age = 28.8; SD = 3.6). For more information on recruitment, inclusion and exclusion criteria, protocols for the data collection and image acquisition for the MR-scans, please see Uğurbil et al., 2013; Van Essen et al., 2012; Van Essen et al., 2013. This study was approved by the regional ethics comity for medical research.

### Image Processing

The current study utilized the minimally pre-processed HCP data (HCP Minimal Pre-processing Pipeline), for more information please see Glasser et al., 2013; HumanConnectomeProject, 2018. All further processing was performed within SPM12 (https://www.fil.ion.ucl.ac.uk/spm/software/spm12), and included smoothing with a 6mm Gaussian kernel and denoising, prior to extracting the time series for the DCM analysis. For denoising the data, a two-step general linear model (GLM) was specified to regress out non-grey matter sources of noise and movement artefacts. Two regions of interest (ROIs, sphere with a radius of 6mm) were specified for each subject, one within the white matter (MNI coordinate: [0 −24 −33]) and one within the cerebrospinal fluid (MNI coordinate: [0 −40 −5]). To control for subject movement in the course of the scanning session, twelve movement parameters were included in the GLM analysis. As the rs-data was acquired in two scanning sessions per individual (Van Essen et al., 2012), the preprocessing steps were performed separately for each of the sessions (Left/Right and Right/Left phase encoding) and for each individual separately.

## DCM Analyses

To study the effect of BP, HCT, BMI and HbA1c on the hemodynamics and effective connectivity in three large scale rs-networks, the rs-fMRI time-series from eight ROIs (sphere with a radius of 6mm) were extracted. See **Table 1** for the coordinates for each region and which network they are a part of. The number of included ROIs were limited to eight due to computational strain.

**Table 1.** Coordinates for Regions of Interest

Coordinates for Regions of Interest
| Resting State Network | Regions | R/L | MNI Coordinates (x, y, z) |
| --- | --- | --- | --- |
| Default Mode Network | PCC | R/L | 0, -52, 26 |
|  | mPFC | R/L | 3, 54, -2 |
|  | LIPC | L | -50, -63, 32 |
|  | RIPC | R | 48, -69, 35 |
| Central Executive Network | DLPFC | R | 45, 16, 45 |
|  | PPC | R | 54, -50, 50 |
| Salience Network | AI | R | 37, 25, -4 |
|  | ACC | R/L | 4, 30, 30 |

*Note.* The coordinates for each ROI, and which rs-network the given regions is considered a part of. Colors correspond the **Figure 1**. MNI = Montreal Neurological Institute; R = right hemisphere; L=left hemisphere. *Abbreviations:* PCC = Posterior Cingulate Cortex; mPFC = Medial Prefrontal Cortex; LIPC = Left Inferior Parietal Cortex; RIPC = Right inferior Parietal Cortex; DLPFC = Dorsolateral Prefrontal Cortex; PPC = Posterior Parietal Cortex; AI = Anterior Insular Cortex; ACC = Anterior Cingulate Cortex.

The time series of each ROI were extracted and subjected to individual csd-DCM analyses, and the csd-DCMs were specified for each rs-fMRI acquisition separately; as each subject underwent two rs-fMRI acquisitions, a total of 1188 acquisitions were included in the analysis. In the current study, the *priors* were turned on for connections between all the included regions. The csd-DCM resulted in parameters of effective connectivity between and within each region (A-matrix) and hemodynamic parameters (*transit time* for each region, *epsilon* and *decay*). In addition, csd-values (α- and β-values) and Free Energy parameters were extracted.

The effective connectivity (A-matrix) refers to the correlation in the frequency distribution of the BOLD-signal between brain regions. As the activity in one region is being modelled as a function of the frequency distribution in another region, it indicates the causal interference one brain region makes on another (Friston et al., 2014; Zeidman et al., 2019a). The hemodynamic parameters are essentially descriptions of the Balloon Model; the *transit time* for each predefined region (resting CBV divided by resting CBF, as a measure of BOLD-signal dynamics), *decay* as the global parameter of the BOLD-signal (the reduction of the BOLD-signal, which relates to the relaxation of the smooth musculature of the arterioles), and *epsilon* as the neuronal efficacy (the relationship between CBF and neuronal activity, reflecting an increase in relative CBF expressed as a number of transients per second (Friston, Mechelli, Turner & Price, 2000; Zeidman et al., 2019a). The spectral density values, expressed as α- and β-values, reflect the amplitudes and exponents of the spectral density of the neuronal fluctuations, respectively. Further, model evidence, expressed as “Free Energy” is calculated. If the model evidence increases by including variables in the model, it indicates that more of the model is explained when the given variables are included (Friston et al., 2014).

### Statistical analyses

To test the hypotheses of between-subject variation in the DCM parameters, a hierarchical linear regression analyses was conducted. As some relationship between the independent variables is assumed, a hierarchical linear regression analysis was *only* conducted on the independent variables (BP, HCT, BMI and HbA1c) and the dependent variables (DCM parameters) that showed a significant correlation (Pearson two-tailed *p* < .05). If more than one independent variable correlated significantly with the same dependent variable, both were added to the same regression model, while controlling for the effect of gender. For the full models (including the independent variable(s) and gender) it was examined which of the variables that made a significant unique contribution to explaining the variance in the dependent variable, and whether the full model significantly predicted the dependent variable. To investigate the relative contribution of each factor independently, the standardized coefficient Standardized Beta of the coefficients t-test were examined, as well as R^2^ Change for the F-test of the full models. Extreme outliers, defined as >3.0 interquartile from the mean were removed. The alpha value was set to 0.05, and the confidence interval was set to 95%. The effective connectivity parameters LIPC to ACC, RIPC, and PPC to mPFC did not meet the assumptions and were therefore not included in the final regression analysis.

In addition, a Kruskal-Wallis H Test of between group variance was conducted, comparing three BMI groups on the DCM parameters. A non-parametric test was chosen over a parametric test of group comparison, as the groups were not normally distributed. The BMI variable was split into three groups; normal weight (BMI 19-24, *n* = 262), overweight (BMI 25-29, *n* = 202) and obese (BMI >30, *n* = 117), for comparison. The χ^2^-distribution was defined by the *degrees of freedom* (K-1), which in this case was 2. In the cases of significant group differences, a pairwise comparison with Bonferroni corrections was used to investigate which groups significantly differed (*p* < .05). The Mean Rank Value was used to determine which of the groups showed a higher rank order, and effect sizes were calculated (*r* = z/√*n*+*n*). (See Supplementary Figure 1 for the frequency distribution of the groups across BMI scores/groups).

## Results

### Descriptive Statistics

A two-tailed Pearson correlation analysis was conducted with the independent variables (confidence interval = 95%). It revealed a significant relationship (*p* < .05) between systolic BP and HCT (*r* = .105, *p* = .015) and systolic BP and BMI (*r* = .335, *p* = .000). There was also a significant relationship between diastolic BP and HCT (*r* = .085, *p* = .048) and diastolic BP and BMI (*r* = .291, *p* = .000). Systolic and diastolic BP had the strongest significant covariation (*r* = .680, *p* = .000). (See Supplementary Table 1 for the full correlation matrix).

### Hemodynamic parameters

For mPFC *transit time*, adding BMI and diastolic BP explained significantly more of the variance, both variables made significant unique contributions to the model, and the full model was significant; diastolic BP and BMI together explained 2.8% of the variance. For AI *transit time*, adding diastolic BP to the model made a significant, unique, contribution to explaining the variance, and the full model was significant; explaining 1.3% of the variance.

For the remaining hemodynamic parameters, namely *decay* and *epsilon*, diastolic BP and systolic BP made a significant contribution to the model for *epsilon*, with diastolic BP making a significant unique contribution to explaining the variance. The full model was significant; explaining 1.7% of the variance. See **Table 2.A** for the regression table for these parameters.

**Table 2.**
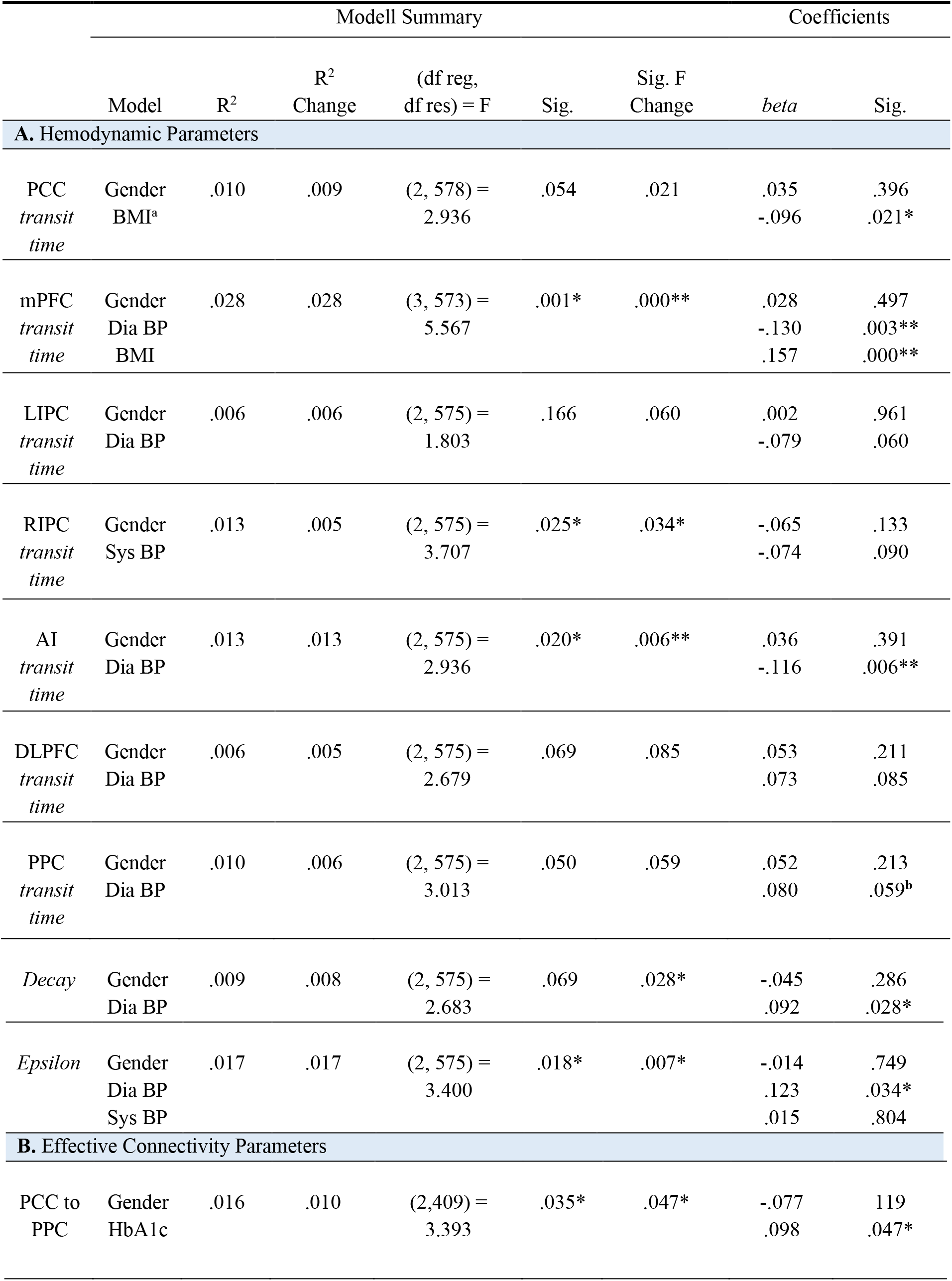
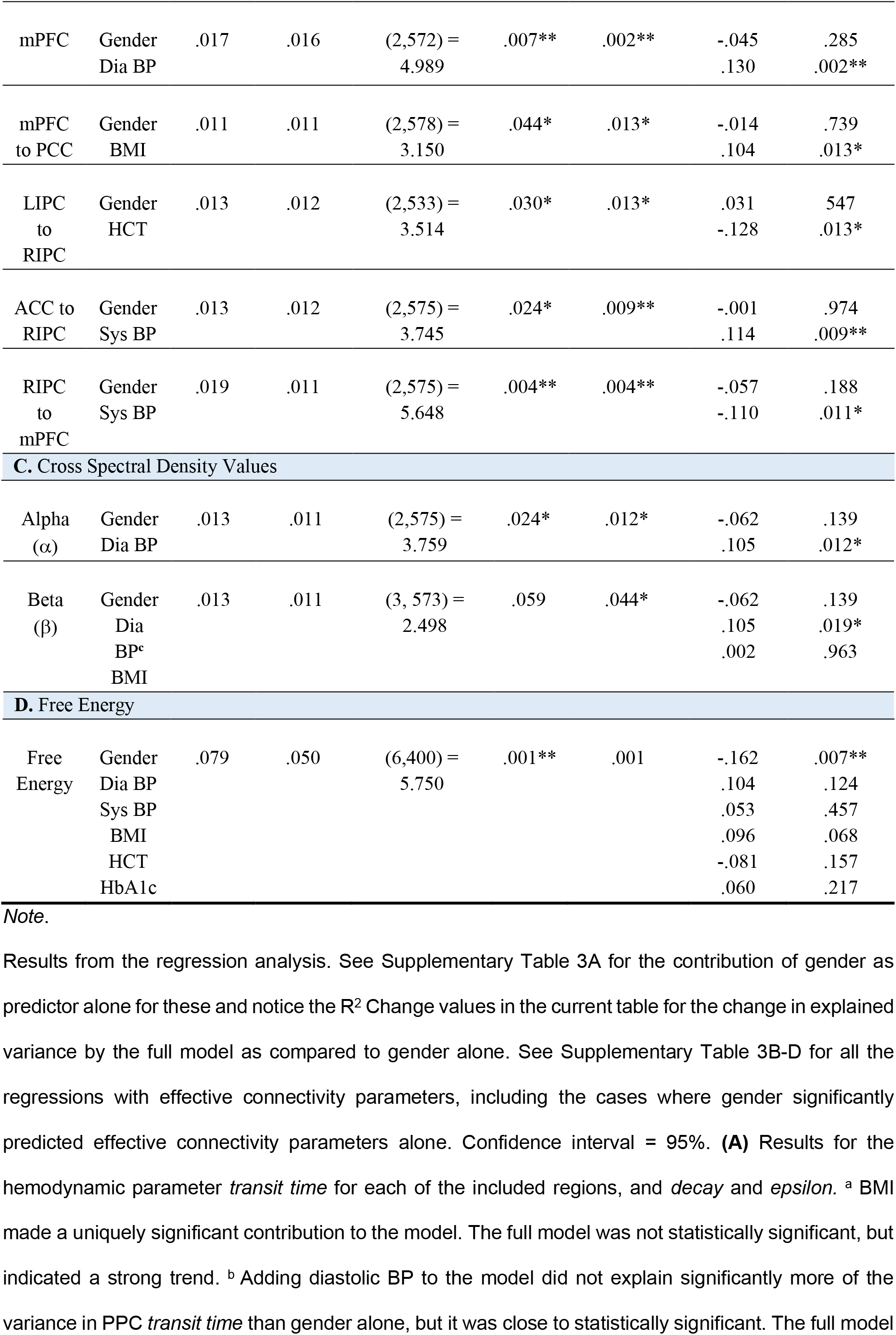
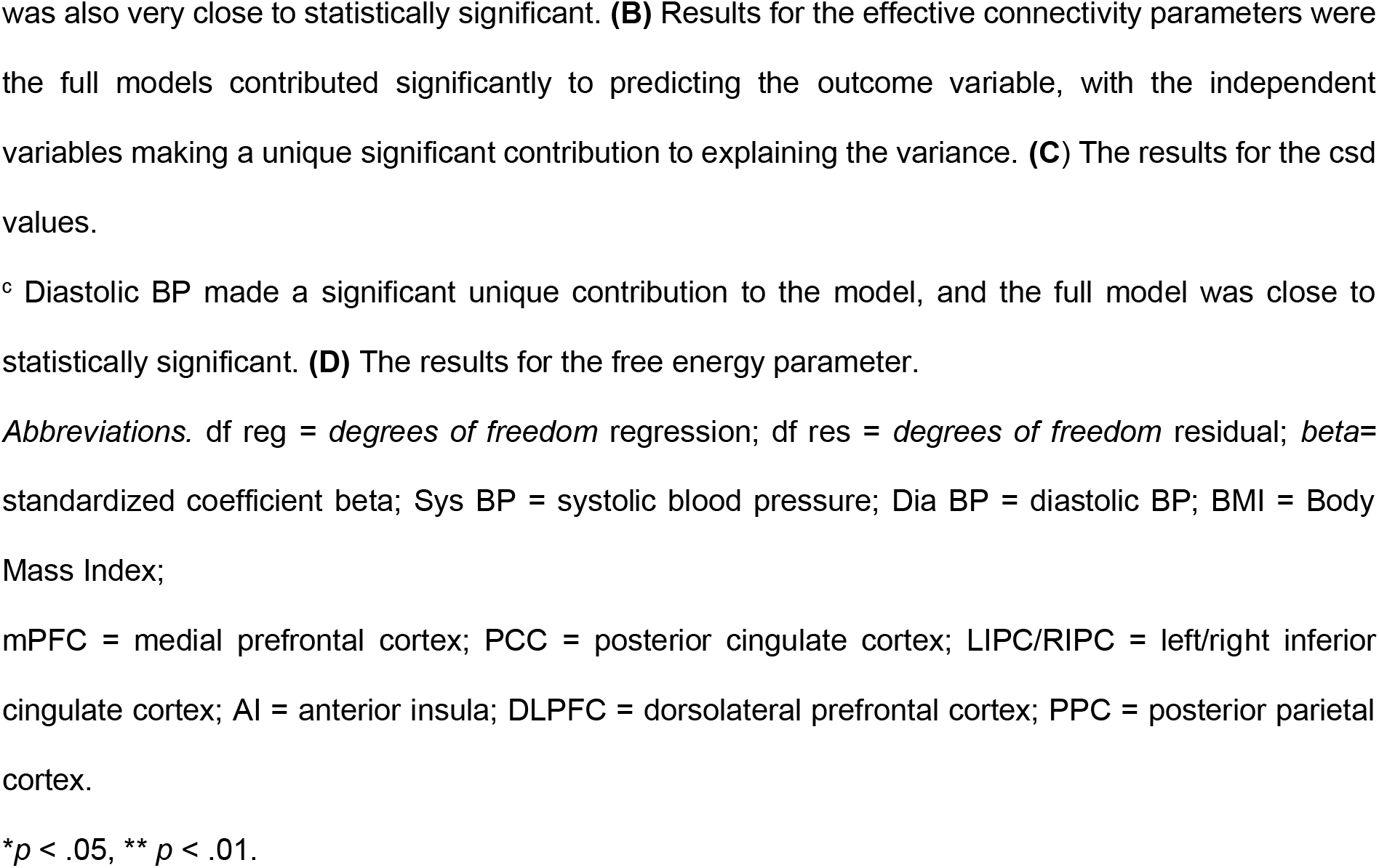
Regression Table for Significant Results

Regression Table for Significant Results
|  |  | Modell Summary |  |  |  |  | Coefficients |  |
| --- | --- | --- | --- | --- | --- | --- | --- | --- |
|  | Model | R<sup>2</sup> | R<sup>2</sup><br>Change | (df reg,<br>df res) = F | Sig. | Sig. F<br>Change | *beta* | Sig. |
| **A. Hemodynamic Parameters** |  |  |  |  |  |  |  |  |
| PCC<br>*transit*<br>*time* | Gender<br>BMI<sup>a</sup> | .010 | .009 | (2, 578) =<br>2.936 | .054 | .021 | .035<br>-.096 | .396<br>.021* |
| mPFC<br>*transit*<br>*time* | Gender<br>Dia BP<br>BMI | .028 | .028 | (3, 573) =<br>5.567 | .001* | .000** | .028<br>-.130<br>.157 | .497<br>.003**<br>.000** |
| LIPC<br>*transit*<br>*time* | Gender<br>Dia BP | .006 | .006 | (2, 575) =<br>1.803 | .166 | .060 | .002<br>-.079 | .961<br>.060 |
| RIPC<br>*transit*<br>*time* | Gender<br>Sys BP | .013 | .005 | (2, 575) =<br>3.707 | .025* | .034* | -.065<br>-.074 | .133<br>.090 |
| AI<br>*transit*<br>*time* | Gender<br>Dia BP | .013 | .013 | (2, 575) =<br>2.936 | .020* | .006** | .036<br>-.116 | .391<br>.006** |
| DLPFC<br>*transit*<br>*time* | Gender<br>Dia BP | .006 | .005 | (2, 575) =<br>2.679 | .069 | .085 | .053<br>.073 | .211<br>.085 |
| PPC<br>*transit*<br>*time* | Gender<br>Dia BP | .010 | .006 | (2, 575) =<br>3.013 | .050 | .059 | .052<br>.080 | .213<br>.059<sup>b</sup> |
| *Decay* | Gender<br>Dia BP | .009 | .008 | (2, 575) =<br>2.683 | .069 | .028* | -.045<br>.092 | .286<br>.028* |
| *Epsilon* | Gender<br>Dia BP<br>Sys BP | .017 | .017 | (2, 575) =<br>3.400 | .018* | .007* | -.014<br>.123<br>.015 | .749<br>.034*<br>.804 |
| **B. Effective Connectivity Parameters** |  |  |  |  |  |  |  |  |
| PCC to<br>PPC | Gender<br>HbA1c | .016 | .010 | (2,409) =<br>3.393 | .035* | .047* | -.077<br>.098 | .119<br>.047* |
| mPFC | Gender<br>Dia BP | .017 | .016 | (2,572) =<br>4.989 | .007** | .002** | -.045<br>.130 | .285<br>.002** |
| mPFC<br>to PCC | Gender<br>BMI | .011 | .011 | (2,578) =<br>3.150 | .044* | .013* | -.014<br>.104 | .739<br>.013* |
| LIPC<br>to<br>RIPC | Gender<br>HCT | .013 | .012 | (2,533) =<br>3.514 | .030* | .013* | .031<br>-.128 | .547<br>.013* |
| ACC to<br>RIPC | Gender<br>Sys BP | .013 | .012 | (2,575) =<br>3.745 | .024* | .009** | -.001<br>.114 | .974<br>.009** |
| RIPC<br>to<br>mPFC | Gender<br>Sys BP | .019 | .011 | (2,575) =<br>5.648 | .004** | .004** | -.057<br>-.110 | .188<br>.011* |
| **C. Cross Spectral Density Values** |  |  |  |  |  |  |  |  |
| Alpha<br>($\alpha$) | Gender<br>Dia BP | .013 | .011 | (2,575) =<br>3.759 | .024* | .012* | -.062<br>.105 | .139<br>.012* |
| Beta<br>($\beta$) | Gender<br>Dia<br>BP<sup>c</sup><br>BMI | .013 | .011 | (3, 573) =<br>2.498 | .059 | .044* | -.062<br>.105<br>.002 | .139<br>.019*<br>.963 |
| **D. Free Energy** |  |  |  |  |  |  |  |  |
| Free<br>Energy | Gender<br>Dia BP<br>Sys BP<br>BMI<br>HCT<br>HbA1c | .079 | .050 | (6,400) =<br>5.750 | .001** | .001 | -.162<br>.104<br>.053<br>.096<br>-.081<br>.060 | .007**<br>.124<br>.457<br>.068<br>.157<br>.217 |

*Note.*
was also very close to statistically significant. **(B)** Results for the effective connectivity parameters were the full models contributed significantly to predicting the outcome variable, with the independent variables making a unique significant contribution to explaining the variance. **(C)** The results for the csd values.
<sup>c</sup> Diastolic BP made a significant unique contribution to the model, and the full model was close to statistically significant. **(D)** The results for the free energy parameter.
*Abbreviations.* df reg = *degrees of freedom* regression; df res = *degrees of freedom* residual; *beta*= standardized coefficient beta; Sys BP = systolic blood pressure; Dia BP = diastolic BP; BMI = Body Mass Index;
mPFC = medial prefrontal cortex; PCC = posterior cingulate cortex; LIPC/RIPC = left/right inferior cingulate cortex; AI = anterior insula; DLPFC = dorsolateral prefrontal cortex; PPC = posterior parietal cortex.
\* $p < .05$ , \*\* $p < .01$ .

The only hemodynamic parameter that differed between BMI groups was PCC *transit time* (normal weight *n* = 262, overweight *n* = 202, obese *n* = 117), χ^2^ (*df* = 2, *n* = 581) = 8.718, *p* = .013. After the pairwise comparison with Bonferroni corrections, the significant difference between the groups were found to be between the overweight and the obese group, with the overweight group having a higher mean rank value (*mean rank* = 314) than the obese group (*mean rank* = 258), *p* = .009, *r* = 0.166. (See Supplementary Table 2 for the Kruskal-Wallis H tests with all hemodynamic parameters).

### Effective connectivity parameters

The regressions with effective connectivity parameters as outcome variables revealed some of the independent variables to contribute significantly to explaining more of the variance in the given parameter than gender, and that one or more of the independent variables contributed significantly to this tendency, with the full model (including gender) being significant (*p* = < .05). HbA1c made a significant contribution to explaining the variance in PCC to PPC, explaining 1.0% of the variance. Diastolic BP significantly contributed to explain the variance in mPFC, and explaining 1.6% of the variance. BMI made a significant contribution to explaining the variance in mPFC to PCC, explaining 1.1% of the variance. HCT made a significant contribution to explaining the variance in LIPC to RIPC, explaining 1.2% of the variance. Systolic BP made a significant contribution to explaining the variance in ACC to RIPC, explaining 1.2% of the variance. Systolic BP made a significant contribution to explaining the variance in RIPC to mPFC, and the full model was significant, with systolic BP explaining 1.1% of the variance **(see table 2.B).**

The Kruskal-Wallis H Test revealed significant BMI group differences, which can be seen in **Table 3**, and **Figure 1**. After the pairwise comparison with Bonferroni corrections, the overweight group and the obese group significantly differed in the effective connectivity from PCC to mPFC, from PCC to RIPC, and within RIPC, with the overweight group having the highest mean rank value. The normal weight group and the overweight group significantly differed on effective connectivity from LIPC to PCC, from LIPC to mPFC, from LIPC to RIPC, from AI to mPFC and within RIPC. The normal weight group had the highest mean rank value for all the connections except for from AI to mPFC, where the overweight group had the highest mean rank value. The normal weight and the obese group significantly differed in effective connectivity from PCC to DLPFC and from mPFC to RIPC. For PCC to DLPFC the normal weight group had the highest mean rank value, indicating stronger effective connectivity, whereas the obese group had a higher mean rank value from mPFC to RIPC. As can be seen from **Table 3,** the effect sizes are low for all the significant different groups (Cohen, 1988). See Supplementary Table 4 for the non-significant between-group results.

**Figure 1.**
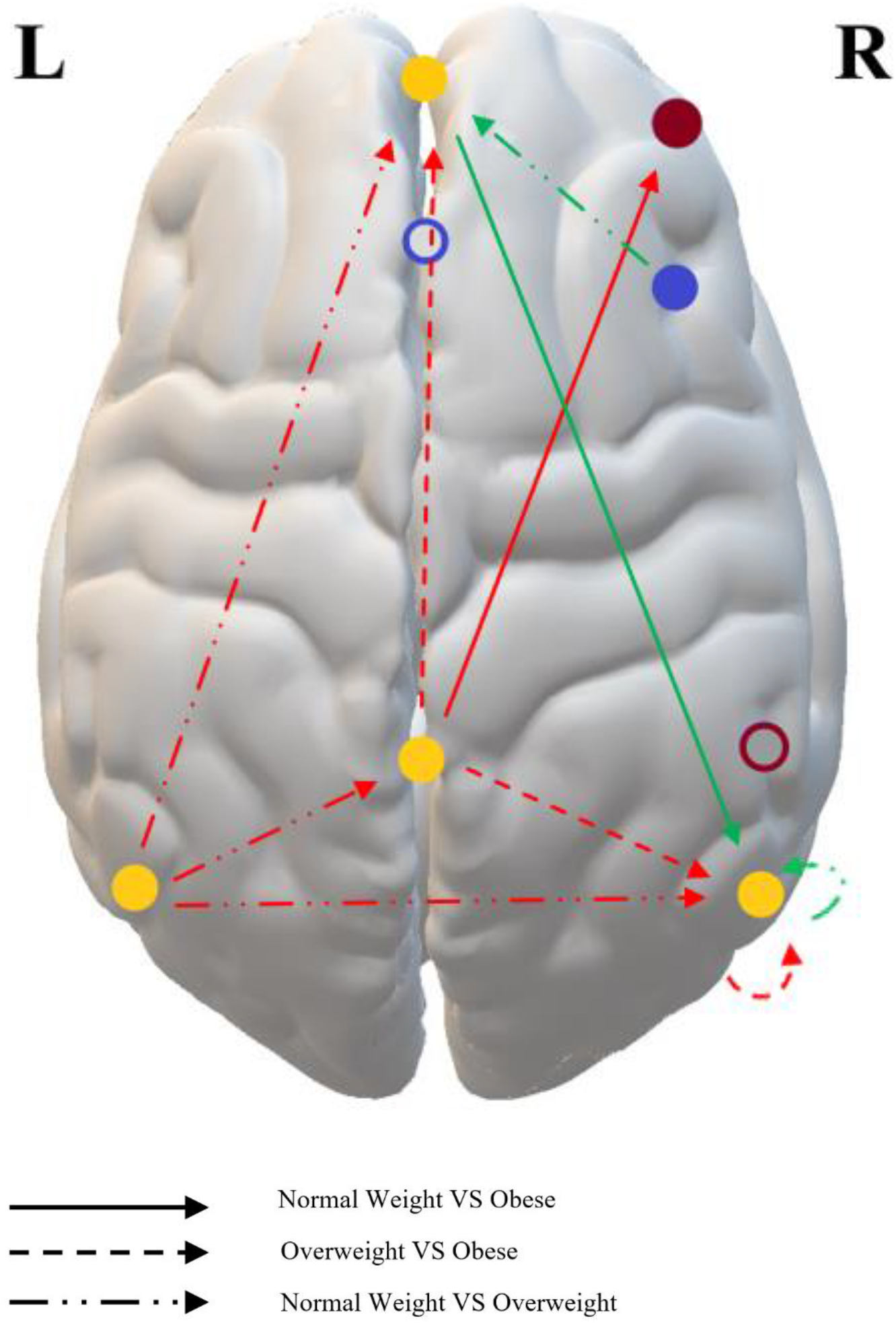
Significant Differences between BMI-groups in Effective Connectivity Parameters *Note*. Illustration of the significant group-differences of the Kruskal-Wallis H Test for the effective connectivity parameters. Please see Table 1 for the exact MNI coordinates for each region. The arrows indicate which groups are being compared, and the directionality of the connection. Based on the mean rank value of the given groups in the comparison, it was determined whether the connection was strengthened (green arrow) or weakened (red arrow) with increased BMI. The color of the solid circles indicates the networks in which the regions showed sig. group differences; yellow being DMN (mPFC, PCC, LIPC, RIPC), dark red being CEN (DLPFC) and blue being SN (AI). The white circles with a colored border are the regions that did not exhibit significant groups differences; PPC (dark red circle, as it is a part of CEN) and ACC (blue circle, as it is a part of SN). As can be seen from the figure, all of the affected connections are within DMN, or between DMN and CEN or SN. mPFC = medial prefrontal cortex; LIPC/RIPC = left/right inferior parietal cortex; PCC = posterior cingulate cortex; DLPFC = dorsolateral prefrontal cortex; PPC = posterior parietal cortex, AI = anterior insula; ACC = anterior cingulate cortex. Please see Table 3 for the full analysis.

**Table 3.**
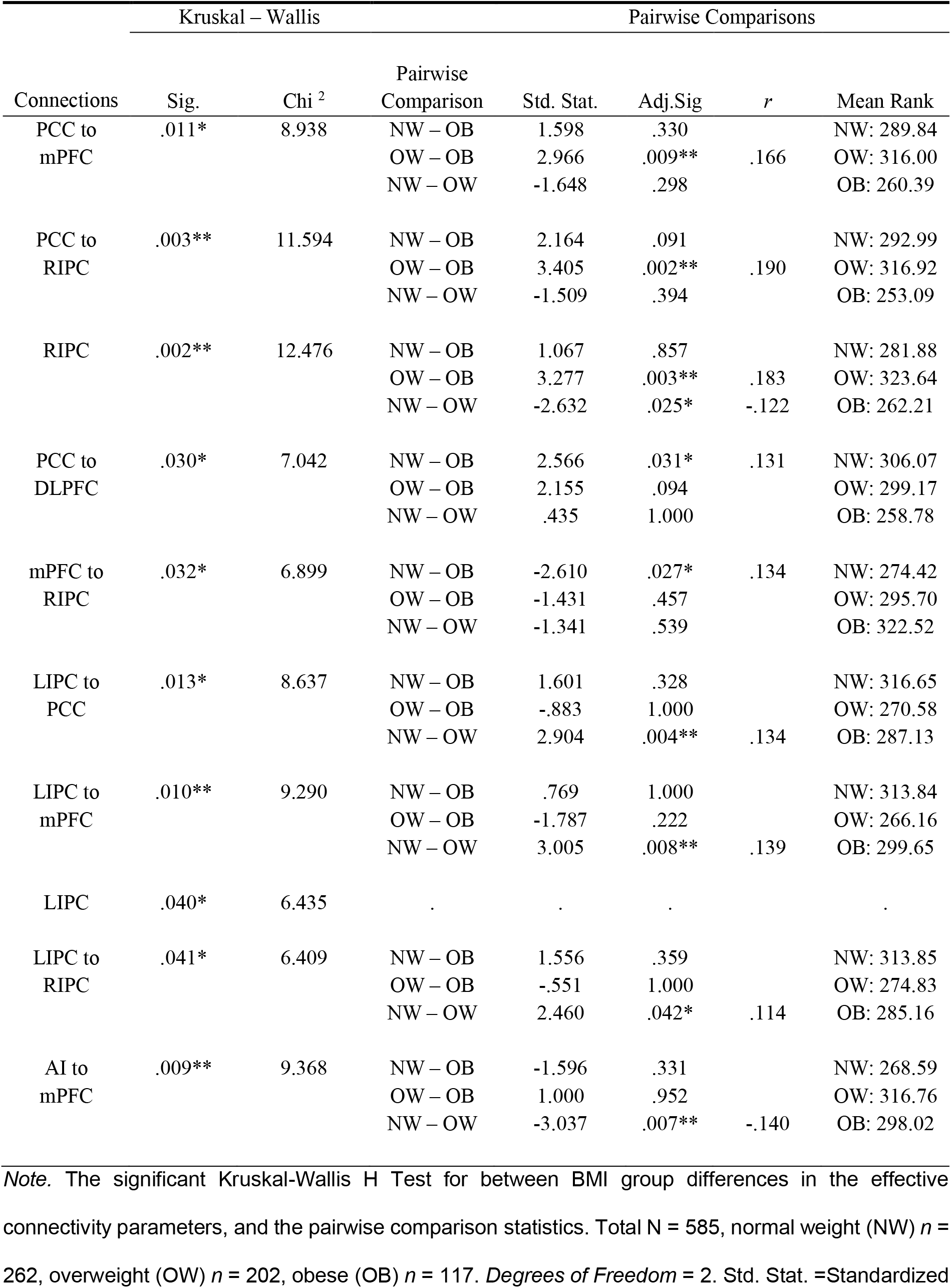
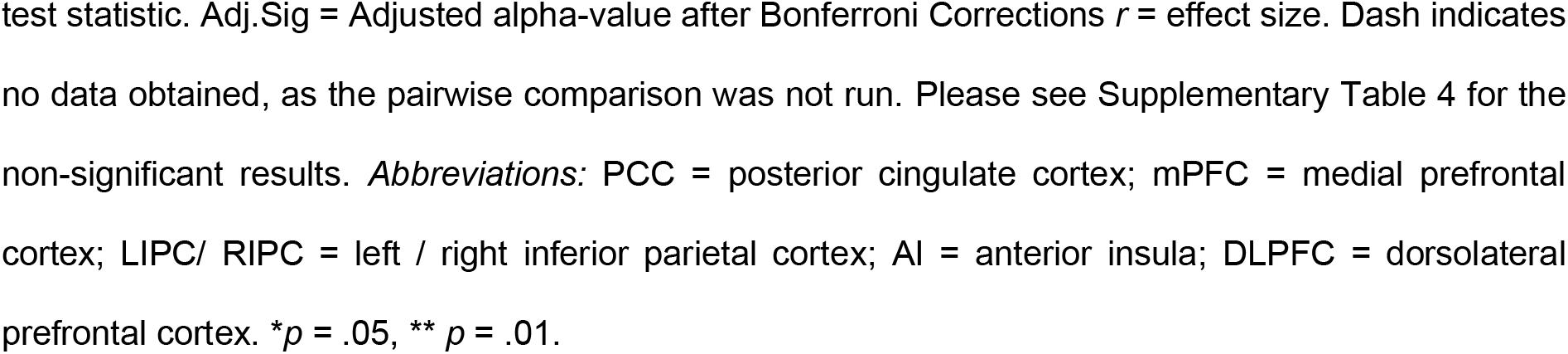
Sig. Kruskal-Wallis H Tests with Effective Connectivity Parameters, with sig. Group Differences

### Cross spectral density parameters

Diastolic BP made a significant contribution to explaining the variance in the α-value, explaining 1.1% of the variance (**see table 2.C**). The Kruskal-Wallis H test revealed no significant group differences on the csd parameters α-value or ß-value between the BMI groups; α-value (normal weight *n* = 262, overweight *n* = 202, obese *n* = 117), χ^2^ (*df* = 2, *n* = 581) = .439, *p* = .803, and ß-value (normal weight *n* = 262, overweight *n* = 202, obese *n* = 117), χ^2^ (*df* = 2, *n* = 581) = 2.644, *p* = .267.

### Free energy

For the free energy parameter, the full regression model explained significantly more than gender alone. However, gender made the only significantly unique contribution to the full model. The full model explains around 7.9% of the variance in Free Energy, and BP, HCT, BMI and HbA1c accounted for 5.0% of the explained variance (see **Table 2.D**).

## Discussion

Several of the hypotheses, regarding variability in hemodynamic and effective connectivity parameters, were supported. The results of the regression analyses imply that between-subject variance in the hemodynamic parameters of rs-fMRI is predicted by diastolic BP and BMI. Further, the variance in the effective connectivity within and between rs-networks is predicted by diastolic and systolic BP, HCT, BMI and HbA1c. The between-group comparison for BMI gives insight into hemodynamic and effective connectivity differences between BMI groups. Although, the effect sizes were quite small, but as can be seen from the regression analysis with Free Energy, including the independent variables increases the explained model evidence; collectively explaining around 5 % of the variance.

### Hemodynamic Parameters – Effects of Blood Pressure and BMI

When including BP in the full regression models, it was found that an increase in diastolic BP has an impact on the shape and amplitude of the BOLD-signal, with a globally increased amplitude of the BOLD-signal, as reflected by the global parameter *epsilon* and the csd-parameter α, and a locally shortening in the regions of mPFC and AI, as indicated by an inverse relationship with the local parameter *transit time*. The shortened *transit time* confirm our hypothesis for mPFC and AI, and the hemodynamic signal appear to be altered in all the regions of interest. The results indicate that diastolic BP affects the rate at which oHb is being utilized in the capillaries and dHb is transported to the veins; essentially weakening the response from the venous compartments in terms of the Balloon model proposed by Buxton et al. (1998). One explanation of this tendency might be related to stiffer venules and veins, which could increase diastolic BP, and prevent the Balloon effect (Buxton, 2012; Buxton et al., 1998). The explanation of the observed variance is however probably multifaceted, as the dynamics of the BOLD-signal is complex and generally poorly understood (Arthurs & Boniface, 2002; Ekstrom, 2010; Handwerker, Gonzalez-Castillo, D’esposito & Bandettini, 2012; Logothetis & Wandell, 2004). For example, we observed a strong trend for diastolic BP and PPC *transit time* in the opposite direction of the results for diastolic BP and AI/mPFC.

Increased BMI was found to prolong the BOLD-signal in mPFC while the shape/amplitude remain the same, in our regression model. However, the BOLD-signal was shortened in PCC with high BMI in the group comparison analysis (overweight group vs obese group). A similar trend was found for PCC in the regression model, though only close to significant. Higher BMI is associated with decreased CBV, which is thought to alter the Balloon effect in terms of making the BOLD-signal shorter (Buxton et al., 1998; Lemmens, Bernstein & Brodsky, 2006). As the results from the regression analysis and the group-comparison differ somewhat, in terms of the of the results’ direction, it might be the case that the hemodynamic response in different brain regions are affected differently by increased BMI.

The effect BMI and BP have on *transit time* is not evident in all the regions within the examined rs-networks, but are limited to mPFC and AI for diastolic BP, and mPFC and PCC for BMI. mPFC and AI are situated close to two of the big arteries of the brain, namely arteria cerebri anterior and media, which might be one explanation of why these regions appear to be sensitive to differences in hemodynamics. These results are highly intriguing as mPFC and AI are the main hubs of DMN and SN, respectively, and, interestingly, that the BOLD signal of mPFC is affected by both diastolic BP and BMI. Hence, the variability in diastolic BP and BMI might have affected the results of previously conducted functional connectivity studies on DMN and SN, while actually being attributable to hemodynamic variability (Rangaprakash, Wu, Marinazzo, Hu & Deshpande, 2018).

The amplitude of the low frequency fluctuations (ALFFs) of the rs-fMRI BOLD-signal and the amplitude of the frequency of cardiovascular fluctuations are within the same range; below 0.10 Hz for the BOLD-signal and around 0.08 Hz for the cardiovascular fluctuations (Zhu, Tarumi, Khan & Zhang, 2015). The results of the current study are in line with previous results finding the BOLD-signal to be highly coupled with cardiovascular fluctuations and CBF globally, and with regions within DMN showing the strongest significant coupling (Kobuch, Macefield & Henderson, 2019; Tak, Wang, Polimeni, Yan & Chen, 2014; Whittaker, Driver, Venzi, Bright & Murphy, 2019). While the amount of explained variance by diastolic BP and BMI alone is fairly low, when considering all of the endogenous and exogenous factors that have been found to affect connectivity of rs-networks and the BOLD-signal, these components might be contributing factors to the overall low replicability and variability in rs-fMRI studies.

### Effective Connectivity

For BP, there was a positive relationship between the diastolic BP and the “self-inhibitory connection of” mPFC, which indicates that mPFC is more inhibited with increased diastolic BP. Further, the systolic BP displayed a negative relationship with connections within DMN, and a positive relationship for the connectivity from SN to DMN. The relationships between BP and some of the connections might be related to previous research, which indicated altered connectivity with hypertension in combination with cognitive impairments (Bu et al., 2018).

In the regression analysis, increased BMI predicted a strengthening in one of the connections within DMN (mPFC to PCC), which is not in line with our hypothesis. By contrast, the comparison of the different BMI groups did to a large extent confirm our hypothesis. In short, the results indicate an internal weakening of the connectivity within DMN with increased BMI, as well as stronger connectivity from SN to DMN. The results also indicate that the normal weight group, compared to the obese group, have stronger connections from DMN to CEN. The only connection within DMN that was stronger with increased BMI in the group-comparison, was from mPFC to RIPC, for the obese compared to the normal weight group. It should be noted that, when taking the mean rank of all the groups into consideration, a non-linear relationship appears for some of the effective connectivity parameters. This might be why the results of the group comparison differ somewhat from the regression analysis, as it assumes a linear relationship. It might also be why previous research has indicated a linear relationship between higher BMI and changes in rs-connectivity, as most of the studies compare only two groups. This relationship could be further investigated in future studies.

As suggested in previous studies, the result might be reflecting poorly regulated eating behavior, as networks involved in balancing sensory-driven and internally guided behavior are affected (Doucet et al., 2017; Sadler et al., 2018). The current study contributes to a deeper understanding of the interactions between these networks, as effective connectivity not functional connectivity is investigated. It is for example interesting to notice that the strengthened connection between AI and mPFC with increased BMI, namely between SN and DMN, is in fact in the direction *from* SN *to* DMN. In addition, connections from LIPC to several regions in DMN appear to be weaker with increased BMI, which might indicate that LIPC mediates the weakened connectivity in the other regions of DMN (PCC, mPFC and RIPC).

As expected, the internal connectivity of DMN was weakened by increased HCT. However, previous studies have indicated that HCT accounts for a weakening of the internal cohesiveness of *all* of the rs-networks included in this study (Yang et al., 2015). As the assumptions of regression was only met for LIPC to RIPC, it was the only conducted analysis with HCT. However, one might consider the correlation analysis as indicative of minimal relationship between HCT and the effective connectivity of the rs-networks.

HbA1c exhibited a positive relationship with the effective connectivity from DMN to SN which partly confirms the a-prior hypothesis. A weakening of the internal connectivity in CEN and SN was however not observed. Previous research has indicated altered connectivity in AI and its connections, but in our case the assumptions of regression was not being met for many of the analysis that included HbA1c as predictor and connections with AI (the only one included in a regression model was AI to ACC). HbA1c did not contribute to explaining the variance in this case. However, the previous results have mostly compared subjects with pre-diabetes or diabetes, as opposed to the normal scores of the current study (Liu et al., 2017; Sadler et al., 2019; Yang et al., 2016).

### General discussion

Seen together, the results indicate that all the factors included in this study can affect the underlying connectivity of the resting human brain, and that diastolic BP and BMI specifically can affect the measured BOLD-signal. As the effect sizes and the percent of explained variance are low, these factors are likely to not cause a large degree of variability in the results independently. However, as implied by Free Energy, they might collectively contribute to unreliable results. Further, the current study adds to the growing body of research on rs-fMRI variability, as the results support the notion that variability in the BOLD-signal and connectivity of rs-networks is to be found even within a fairly healthy population. Taking BP, HCT, BMI and HbA1c, into account when studying rs-networks might thus be advantageous for future studies, to increase the studies reproducibility. Further, some of the benefits of using DCM are highlighted with the current study. Utilizing the DCM technique enables future studies to assess whether their results arise from neuronal changes, as well as the directionality of these changes, or if the changes merely represent variation in cerebral hemodynamics.

As the results of this study indicate that the included endogenous factors are associated with changes in the underlying connectivity *and* hemodynamic variability of rs-networks, it adds to the number of studies that undermine the notion of rs-networks as being stable between subjects in terms of connectivity, as well as being susceptible to hemodynamic variation caused by BP and BMI. Interestingly, DMN seems to be the most inherently unstable of the rs-networks, as it is susceptive to vary with the endogenous factors in both hemodynamic and effective connectivity parameters. Seen together with results from other studies, the wide range of potential sources of variability might be one of the reasons why it has been proven a challenge to come to a unified view on the function of DMN. In addition, the results implicate that DMN might not inhabit the between-subject stability which it is ascribed when studying clinical deviations (Greicius, Srivastava, Reiss & Menon, 2004; Mevel, Chételat, Eustache & Desgranges, 2011).

### Limitations of the Current Study

Alternatively to the current approach, might be the use of Parametric Empirical Bayes (PEB) (Zeidman et al., 2019b), which could not be implemented here for technical reasons. There are some advantages of using PEB with csd-DCM data as compared to other GLMs, and it can therefore be considered a limitation of this study that a more traditional GLM was utilized for the analyses of between-subject effects on the DCM parameters. Specifically, many of the analyses were not performed with HCT and HbA1c as independent variables, as the assumptions of the regression analysis prevented it. Even for the significant correlations, which “allowed” the variables to be entered into the regression analysis, the relationship between the variables were quite low (around .10). Using PEB instead would allow for all the scores on all the independent variables to be analyzed with the DCM parameters, to examine their effect.

## Conclusion

The low cost, short scanning times and absence of task, which initially made rs-fMRI an intriguing, useful and popular tool for neuroimaging, comes with a price. Evidently, the rs-networks are susceptible to a variety of exogenous and endogenous factors, which makes the results of the studies unreliable and as a consequence challenging to replicate. This tendency adds to the difficulty of studying and understanding the function of rs-networks like DMN and using alterations in rs-networks as a clinical marker of disease. Seen together with previous studies and results, the current study is an indicator of the nature of DMN as inheritably unstable, in which case it is unsurprising that its function seems to “slip through the fingers” of the researcher. If rs-networks are to be truly understood, studies should at least take the effect of BP and BMI into account, to ensure more reliable results for the future. By doing so, researcher would also take a step in the right direction in terms of the ongoing replication crisis.

## Acknowledgement and Funding

The study was financed by the Research Council of Norway (Project number: 276044: *When default is not default: Solutions to the replication crisis and beyond*). Data were provided by the Human Connectome Project, WU-Minn Consortium (Principal Investigators: David Van Essen and Kamil Ugurbil; 1U54MH091657) funded by the 16 NIH Institutes and Centers that support the NIH Blueprint for Neuroscience Research; and by the McDonnell Center for Systems Neuroscience at Washington University.

## Notes

https://www.humanconnectome.org/study/hcp-young-adult

### Competing Interest Statement

The authors have declared no competing interest.

## Referances

Agcaoglu, O., Miller, R., Mayer, A. R., Hugdahl, K. & Calhoun, V. D. (2015). Lateralization of resting state networks and relationship to age and gender. NeuroImage, 104(1), 310–325. doi:10.1016/j.neuroimage.2014.09.001

Allen, E. A., Erhardt, E. B., Damaraju, E., Gruner, W., Segall, J. M., Silva, R. F.,… Kalyanam, R. (2011). A baseline for the multivariate comparison of resting-state networks. Frontiers in systems neuroscience, 5, 1–23. doi:10.3389/fnsys.2011.00002

Arthurs, O. J. & Boniface, S. (2002). How well do we understand the neural origins of the fMRI BOLD signal? TRENDS in Neurosciences, 25(1), 27–31.

Biswal, B., Zerrin Yetkin, F., Haughton, V. M. & Hyde, J. S. (1995). Functional connectivity in the motor cortex of resting human brain using echo-planar MRI. Magnetic resonance in medicine, 34(4), 537–541. doi:10.1002/mrm.1910340409

Bressler, S. L. & Menon, V. (2010). Large-scale brain networks in cognition: emerging methods and principles. Trends in cognitive sciences, 14(6), 277–290. doi:10.1016/j.tics.2010.04.004

Bu, L., Huo, C., Xu, G., Liu, Y., Li, Z., Fan, Y. & Li, J. (2018). Alteration in brain functional and effective connectivity in subjects with hypertension. Frontiers in physiology, 9, 1–12. doi:doi: 10.3389/fphys.2018.00669

Buckner, R. L., Andrews-Hanna, J. R. & Schacter, D. L. (2008). The brain’s default network: anatomy, function, and relevance to disease. 1124(1), 1–38. doi:10.1196/annals.1440.011

Buxton, R. B. (2012). Dynamic models of BOLD contrast. NeuroImage, 62(2), 953–961. doi:10.1016/j.neuroimage.2012.01.012

Buxton, R. B., Uludağ, K., Dubowitz, D. J. & Liu, T. T. (2004). Modeling the hemodynamic response to brain activation. NeuroImage, 23, 220–S233. doi:10.1016/j.neuroimage.2004.07.013

Buxton, R. B., Wong, E. C. & Frank, L. R. (1998). Dynamics of blood flow and oxygenation changes during brain activation: the balloon model. Magnetic resonance in medicine, 39(6), 855–864. doi:10.1002/mrm.1910390602

Carnevale, L., Maffei, A., Landolfi, A., Grillea, G., Carnevale, D., & Lembo, G. (2020). Brain functional magnetic resonance imaging highlights altered connections and functional networks in patients with hypertension. Hypertension, 76(5), 1480–1490.

Chao, S.-H., Liao, Y.-T., Chen, V. C.-H., Li, C.-J., McIntyre, R. S., Lee, Y. & Weng, J.-C. (2018). Correlation between brain circuit segregation and obesity. Behavioural brain research, 337(1), 218–227. doi:10.1016/j.bbr.2017.09.017

Choe, A. S., Jones, C. K., Joel, S. E., Muschelli, J., Belegu, V., Caffo, B. S.,… Pekar, J. J. (2015). Reproducibility and temporal structure in weekly resting-state fMRI over a period of 3.5 years. PloS one, 10(10). doi:10.1371/journal.pone.0140134

Cohen, J. W. (1988). Statistical power analysis for the behavioral sciences (2 ed.). Hillsdale, NJ: Lawrence Erlbaum Associates.

Craig, A. D. (2009). How do you feel--now? The anterior insula and human awareness. Nature reviews neuroscience, 10(1), 59–70. doi:10.1038/nrn2555

Curtis, B. J., Williams, P. G., Jones, C. R. & Anderson, J. S. (2016). Sleep duration and resting fMRI functional connectivity: examination of short sleepers with and without perceived daytime dysfunction. Brain and behavior, 6(12). doi:10.1002/brb3.576

Doucet, G. E., Rasgon, N., McEwen, B. S., Micali, N. & Frangou, S. (2017). Elevated body mass index is associated with increased integration and reduced cohesion of sensory-driven and internally guided resting-state functional brain networks. Cerebral Cortex, 28(3), 988–997. doi:10.1093/cercor/bhx008

Ekstrom, A. (2010). How and when the fMRI BOLD signal relates to underlying neural activity: the danger in dissociation. Brain research reviews, 62(2), 233–244. doi:10.1016/j.brainresrev.2009.12.004

Fox, M. D., Snyder, A. Z., Vincent, J. L., Corbetta, M., Van Essen, D. C. & Raichle, M. E. (2005). The human brain is intrinsically organized into dynamic, anticorrelated functional networks. Proceedings of the National Academy of Sciences, 102(27), 9673–9678. doi:10.1073/pnas.0504136102

Friston, K. (2009). Causal modelling and brain connectivity in functional magnetic resonance imaging. PLoS biol, 7(2), https://doi.org/10.1371/journal.pbio.1000033.

Friston, K. J., Kahan, J., Biswal, B. & Razi, A. (2014). A DCM for resting state fMRI. NeuroImage, 94(100), 396–407. doi:10.1016/j.neuroimage.2013.12.009

Friston, K. J., Mechelli, A., Turner, R. & Price, C. J. (2000). Nonlinear responses in fMRI: the Balloon model, Volterra kernels, and other hemodynamics. NeuroImage, 12(4), 466–477. doi:10.1006/nimg.2000.0630

Gauthier, C. J. & Fan, A. P. (2019). BOLD signal physiology: models and applications. NeuroImage, 187(1), 116–127. doi:10.1016/j.neuroimage.2018.03.018

Glasser, M. F., Sotiropoulos, S. N., Wilson, J. A., Coalson, T. S., Fischl, B., Andersson, J. L.,… Polimeni, J. R. (2013). The minimal preprocessing pipelines for the Human Connectome Project. NeuroImage, 80, 105–124. doi:10.1016/j.neuroimage.2013.04.127

Goldstone, A., Mayhew, S. D., Przezdzik, I., Wilson, R. S., Hale, J. R. & Bagshaw, A. P. (2016). Gender specific re-organization of resting-state networks in older age. Frontiers in aging neuroscience, 8, 1–15. doi:10.3389/fnagi.2016.00285

Greicius, M. D., Krasnow, B., Reiss, A. L. & Menon, V. (2003). Functional connectivity in the resting brain: a network analysis of the default mode hypothesis. Proceedings of the National Academy of Sciences, 100(1), 253–258. doi:10.1073/pnas.0135058100

Greicius, M. D., Srivastava, G., Reiss, A. L. & Menon, V. (2004). Default-mode network activity distinguishes Alzheimer’s disease from healthy aging: evidence from functional MRI. Proceedings of the National Academy of Sciences, 101(13), 4637–4642. doi:10.1073/pnas.0308627101

Handwerker, D. A., Gonzalez-Castillo, J., D’esposito, M. & Bandettini, P. A. (2012). The continuing challenge of understanding and modeling hemodynamic variation in fMRI. NeuroImage, 62(2), 1017–1023. doi:10.1016/j.neuroimage.2012.02.015

Harrison, B. J., Pujol, J., Ortiz, H., Fornito, A., Pantelis, C. & Yücel, M. (2008). Modulation of brain resting-state networks by sad mood induction. PloS one, 3(3), 1–12. doi:10.1371/journal.pone.0001794

Hjelmervik, H., Hausmann, M., Osnes, B., Westerhausen, R. & Specht, K. (2014). Resting states are resting traits - An fMRI study of sex differences and menstrual cycle effects in resting state cognitive control networks. PLOS ONE 9 (7): e103492. doi:10.1371/journal.pone.0103492

Hodkinson, D. J., O’daly, O., Zunszain, P. A., Pariante, C. M., Lazurenko, V., Zelaya, F. O.,… Williams, S. C. (2014). Circadian and homeostatic modulation of functional connectivity and regional cerebral blood flow in humans under normal entrained conditions. Journal of Cerebral Blood Flow & Metabolism, 34(9), 1493–1499. doi:10.1038/jcbfm.2014.109

Huang, S. M., Wu, Y. L., Peng, S. L., Peng, H. H., Huang, T. Y., Ho, K. C., & Wang, F. N. (2016). Inter-strain differences in default mode network: a resting state fmri study on spontaneously hypertensive rat and Wistar Kyoto rat. Scientific reports, 6(1), 1–9.

Hugdahl, K., Kazimierczak, K., Beresniewicz, J., Kompus, K., Westerhausen, R., Ersland, L., Grüner, R.,… Specht, K. (2019). Dynamic up- and down-regulation of the default (DMN) and extrinsic (EMN) mode networks during alternating task-on and task-off periods. PLoS ONE, 14(9). https://doi.org/10.1371/journal.pone.0218358

Hugdahl, K., Raichle, M. E., Mitra, A. & Specht, K. (2015). On the existence of a generalized non-specific task-dependent network. Frontiers in Human Neuroscience, 9. doi:10.3389/fnhum.2015.00430

HumanConnectomeProject. (2017). 1200 Subject Data Release. Retrieved from https://www.humanconnectome.org/study/hcp-young-adult/document/1200-subjects-data-release

HumanConnectomeProject. (2018). HCP S1200 Release Referance Manual. Retrieved from https://www.humanconnectome.org/storage/app/media/documentation/s1200/HCP_S1200_Release_Reference_Manual.pdf (30.04.2020)

Kobuch, S., Macefield, V. G. & Henderson, L. A. (2019). Resting regional brain activity and connectivity vary with resting blood pressure but not muscle sympathetic nerve activity in normotensive humans: An exploratory study. Journal of Cerebral Blood Flow & Metabolism, 39(12), 2433–2444. doi:10.1177/0271678X18798442

Koechlin, E. & Summerfield, C. (2007). An information theoretical approach to prefrontal executive function. Trends in cognitive sciences, 11(6), 229–235. doi:10.1016/j.tics.2007.04.005

Lemmens, H. J. M., Bernstein, D. P. & Brodsky, J. B. (2006). Estimating blood volume in obese and morbidly obese patients. Obesity surgery, 16(6), 773–776. doi:10.1381/096089206777346673

Liu, L., Li, W., Zhang, Y., Qin, W., Lu, S. & Zhang, Q. (2017). Weaker functional connectivity strength in patients with type 2 diabetes mellitus. Frontiers in neuroscience, 11(39), 1–11. doi:10.3389/fnins.2017.00390

Logothetis, N. K. & Wandell, B. A. (2004). Interpreting the BOLD signal. Annu. Rev. Physiol., 66(2004), 735–769. doi:10.1146/annurev.physiol.66.082602.092845

Menon, V. & Uddin, L. Q. (2010). Saliency, switching, attention and control: a network model of insula function. Brain Structure and Function, 214(5-6), 655–667. doi:10.1007/s00429-010-0262-0

Mevel, K., Chételat, G., Eustache, F. & Desgranges, B. (2011). The default mode network in healthy aging and Alzheimer’s disease. International journal of Alzheimer’s disease, 2011(special issue), 1–9. doi:10.4061/2011/535816

Raichle, M. E., MacLeod, A. M., Snyder, A. Z., Powers, W. J., Gusnard, D. A. & Shulman, G. L. (2001). A default mode of brain function. Proceedings of the National Academy of Sciences, 98(2), 676–682. doi:10.1073/pnas.98.2.676

Raichle, M. E. & Mintun, M. A. (2006). Brain work and brain imaging. Annu. Rev. Neurosci., 29, 449–476. doi:10.1146/annurev.neuro29.051605.112819

Rangaprakash, D., Wu, G. R., Marinazzo, D., Hu, X. & Deshpande, G. (2018). Hemodynamic response function (HRF) variability confounds resting-state fMRI functional connectivity. Magnetic resonance in medicine, 80(4), 1697–1713. doi:doi.org/10.1002/mrm.27146

Razi, A., Kahan, J., Rees, G., & Friston, K. J. (2015). Construct validation of a DCM for resting state fMRI. Neuroimage, 106, 1–14. https://doi.org/10.1016/j.neuroimage.2014.11.027

Sadler, J. R., Shearrer, G. E. & Burger, K. S. (2018). Body mass variability is represented by distinct functional connectivity patterns. NeuroImage, 181, 55–63. doi:0.1016/j.neuroimage.2018.06.082

Sadler, J. R., Shearrer, G. E. & Burger, K. S. (2019). Alterations in ventral attention network connectivity in individuals with prediabetes. Nutritional neuroscience, 22(4), 1–8. doi:10.1080/1028415X.2019.1609646

Singh, K. D. & Fawcett, I. (2008). Transient and linearly graded deactivation of the human default-mode network by a visual detection task. NeuroImage, 41(1), 100–112. doi:10.1016/j.neuroimage.2008.01.051

Specht, K. (2019). Current challenges in translational and clinical fMRI and future directions. Frontiers in Psychiatry, 10, 1–9. doi:10.3389/fpsyt.2019.00924

Sridharan, D., Levitin, D. J. & Menon, V. (2008). A critical role for the right fronto-insular cortex in switching between central-executive and default-mode networks. Proceedings of the National Academy of Sciences, 105(34), 12569–12574. doi:10.1073/pnas.0800005105

Tak, S., Wang, D. J., Polimeni, J. R., Yan, L. & Chen, J. J. (2014). Dynamic and static contributions of the cerebrovasculature to the resting-state BOLD signal. NeuroImage, 84(1), 672–680. doi:10.1016/j.neuroimage.2013.09.057

Toro, R., Fox, P. T. & Paus, T. (2008). Functional coactivation map of the human brain. Cerebral Cortex, 18(11), 2553–2559. doi:10.1093/cercor/bhn014

Uğurbil, K., Xu, J., Auerbach, E. J., Moeller, S., Vu, A. T., Duarte-Carvajalino, J. M.,… Van de Moortele, P. F. (2013). Pushing spatial and temporal resolution for functional and diffusion MRI in the Human Connectome Project. NeuroImage, 80, 80–104. doi:10.1016/j.neuroimage.2013.05.012

Vaisvilaite, L., Hushagen, V., Grønli, J. & Specht, K. (2020). Time-of-day Effects in resting-state fMRI: changes in Effective Connectivity and BOLD signal. bioRxiv. doi: https://doi.org/10.1101/2020.08.20.258517

Van Essen, D. C., Smith, S. M., Barch, D. M., Behrens, T. E. J., Yacoub, E. & Ugurbil, K. (2013). The WU-Minn Human Connectome Project: An overview. NeuroImage, 80, 62–79. doi:10.1016/j.neuroimage.2013.05.041

Van Essen, D. C., Ugurbil, K., Auerbach, E., Barch, D., Behrens, T., Bucholz, R.,… Curtiss, S. W. (2012). The Human Connectome Project: a data acquisition perspective. NeuroImage, 62(4), 2222–2231. doi:10.1016/j.neuroimage.2012.02.018

Waites, A. B., Stanislavsky, A., Abbott, D. F. & Jackson, G. D. (2005). Effect of prior cognitive state on resting state networks measured with functional connectivity. Human brain mapping, 24(1), 59–68. doi:0.1002/hbm.20069

Whittaker, J. R., Driver, I. D., Venzi, M., Bright, M. G. & Murphy, K. (2019). Cerebral autoregulation evidenced by synchronized low frequency oscillations in blood pressure and resting-state fMRI. Frontiers in neuroscience, 13, 1–12. doi:10.3389/fnins.2019.00433

Yang, S. Q., Xu, Z. P., Zhan, Y. F., Guo, L. Y., Zhang, S., Jiang, R. F.,… Wang, J. Z. (2016). Altered intranetwork and internetwork functional connectivity in type 2 diabetes mellitus with and without cognitive impairment. Scientific reports, 6, 1–11. doi:10.1038/srep32980

Yang, Z., Craddock, R. C. & Milham, M. P. (2015). Impact of hematocrit on measurements of the intrinsic brain. Frontiers in neuroscience, 8, 1–10. doi:10.3389/fnins.2014.00452

Zeidman, P., Jafarian, A., Corbin, N., Seghier, M. L., Razi, A., Price, C. J. & Friston, K. J. (2019a). A guie to group effective connectivity analysis, part 1: first level analysis with DCM for fMRI. NeuroImage, 200, 174–190. doi:10.1016/j.neuroimage.2019.06.031

Zeidman, P., Jafarian, A., Seghier, M. L., Litvak, V., Cagnan, H., Price, C. J. & Friston, K. J. (2019b). A guide to group effective connectivity analysis, part 2: second level analysis with PEB. NeuroImage, 200, 12–25. doi: 10.1016/j.neuroimage.2019.06.032.

Zhu, D. C., Tarumi, T., Khan, M. A. & Zhang, R. (2015). Vascular coupling in resting-state fMRI: evidence from multiple modalities. Journal of Cerebral Blood Flow & Metabolism, 35(12), 1910–1920. doi:10.1038/jcbfm.2015.166

