## Supplementary Material for "Why Blood Pressure and Body Mass Should be Controlled for in Resting-State Functional Magnetic Resonance Imaging Studies"

**Supplementary Materials**

*Supplementary Table 1.*

Correlation Matrix for the Correlation between the Independent Variables

| | | BMI | HbA1c | HCT | Sys BP | Dia BP |
|---|---|---|---|---|---|---|
| BMI | Pearson Correlation | 1 | .050 | .030 | .335 | .291 |
| | Sig. | | .312 | .491 | .000** | .000** |
| | N | 593 | 417 | 547 | 589 | 589 |
| HbA1c | Pearson Correlation | .050 | 1 | -.045 | .063 | .062 |
| | Sig. | .312 | | .363 | .203 | 211 |
| | N | 417 | 417 | 412 | 414 | 414 |
| HCT | Pearson Correlation | .030 | -.045 | 1 | .105* | .085 |
| | Sig. | .491 | .363 | | .015 | .048* |
| | N | 547 | 412 | 547 | 543 | 543 |
| Sys BP | Pearson Correlation | .335 | .063 | .105 | 1 | .680 |
| | Sig. | .000** | .203 | .015* | | .000** |
| | N | 589 | 414 | 543 | 590 | 590 |
| Dia BP | Pearson Correlation | .291 | .062 | .085 | .680 | 1 |
| | Sig. | .000** | .211 | .048* | .000** | |
| | N | 589 | 414 | 543 | 590 | 590 |

*Note.* Correlations between the independent variables BMI, HbA1c, HCT, systolic and diastolic BP. BMI = Body Mass Index, HbA1c = glycated hemoglobin; HCT = hematocrit; Sys BP = systolic blood pressure; Dia BP = diastolic blood pressure.

*Sig. two-tailed $p < .05$, **Sig. two-tailed $p < .01$.

*Supplementary Table 2.*

Kruskal-Wallis H Tests with the Hemodynamic Parameters

| | Kruskal – Wallis H Test | | Pairwise Comparisons | | |
|---|---|---|---|---|---|
| Parameter | Sig. | Chi $^2$ | Pairwise Comparison | Adj.Sig | Mean Rank |
| PCC *transit time* | .013* | 8.718 | OW - OB | .009** | NW: 292.61<br>OW: 314.00<br>OB: 258.66 |
| mPFC *transit time* | .056[a] | 5.750 | . | . | . |
| LIPC *transit time* | .208 | 3.138 | . | . | . |
| RIPC *transit time* | .856 | .310 | . | . | . |
| AI *transit time* | .646 | .875 | . | . | . |
| ACC *transit time* | .230 | 2.938 | . | . | . |
| DLPFC *transit time* | .466 | 1.527 | . | . | . |
| PPC *transit time* | .458 | 1.560 | . | . | . |
| *Decay* | .904 | .201 | . | . | . |
| *Epsilon* | .267 | 2.875 | . | . | . |

*Note.* Kruskal Wallis H Test for between BMI group differences in the hemodynamic parameters. Normal Weight (19-24 kg., n = 262), overweight (25-29 kg., n = 202) and obese (>30 kg., n = 117), Total N = 585, *Degrees of Freedom* = 2. Adj.Sig: Significance level after adjusting the alpha-value for multiple comparisons, with Bonferroni Corrections. Dash indicates no data obtained, as the pairwise comparison was not run. [a] Close to significant group differences for mPFC *transit time*. PCC = posterior cingulate cortex; mPFC = medial prefrontal cortex; LIPC/ RIPC = left / right inferior parietal cortex; AI = anterior insula; ACC = anterior cingulate cortex; DLPFC = dorsolateral prefrontal cortex; PPC = posterior parietal cortex; NW = normal weight; OW = overweight; OB = obese.

**p* = .05, ** *p* = .01.

*Supplementary Table 3.*

Full Regression Table

| | Modell Summary | | | | | | Coefficients | |
|---|---|---|---|---|---|---|---|---|
| | Model | $R^2$ | $R^2$ Change | (df reg, df res) = F | Sig. | Sig. F Change | beta | Sig. |
| **A. Significant results, including gender as predictor alone** | | | | | | | | |
| PCC to PPC | Gender | .007 | | (1,410) = 2.790 | .096 | | | |
| | Gender | .016 | .010 | | .035* | .047* | -.077 | 119 |
| | HbA1c | | | (2,409) = 3.393 | | | .098 | .047* |
| mPFC | Gender | .001 | | (1,575) = .343 | .558 | | | |
| | Gender | .017 | .016 | (2,572) = 4.989 | .007** | .002** | -.045 | .285 |
| | Dia BP | | | | | | .130 | .002** |
| mPFC to PCC | Gender | .000 | | (1,579) = .027 | .870 | | | |
| | Gender | .011 | .011 | (2,578) = 3.150 | .044* | .013* | -.014 | .739 |
| | BMI | | | | | | .104 | .013* |
| LIPC to RIPC | Gender | .001 | | (1,534) = .776 | .379 | | | |
| | Gender | .013 | .012 | (2,533) = 3.514 | .030* | .013* | .031 | 547 |
| | HCT | | | | | | -.128 | .013* |
| ACC to RIPC | Gender | .001 | | (1,576) = .645 | .422 | | | |
| | Gender | .013 | .012 | (2,575) = 3.745 | .024* | .009** | -.001 | .974 |
| | Sys BP | | | | | | .114 | .009** |
| RIPC to mPFC | Gender | .008 | | (1,576) = 4.802 | .029 | | | |
| | Gender | .019 | .011 | | .004** | .004** | -.057 | .188 |
| | Sys BP | | | (2,575) = 5.648 | | | -.110 | .011* |
| Alpha (α) | Gender | .002 | | (1, 576) = 1.205 | .273 | | | |
| | Gender | .013 | .011 | (2,575) = 3.759 | .024* | .012* | -.062 | .139 |
| | Dia BP | | | | | | .105 | .012* |
| Beta (β) | Gender | .002 | | (1, 575) = 1.203 | .273 | | | |
| | Gender | .013 | .011 | (3, 573) = 2.498 | .059 | .044* | -.062 | .139 |

| | | | | | | | | |
|---|---|---|---|---|---|---|---|---|
| | Dia BP[a] | | | | | | .105 | .019* |
| | BMI | | | | | | .002 | .963 |
| Free Energy | Gender | .029 | | (1,405) = 12.063 | .001 | | | |
| | Gender | .079 | .050 | (6,400) = 5.750 | .001** | .001 | -.162 | .007** |
| | Dia BP | | | | | | .104 | .124 |
| | Sys BP | | | | | | .053 | .457 |
| | BMI | | | | | | .096 | .068 |
| | HCT | | | | | | -.081 | .157 |
| | HbA1c | | | | | | .060 | .217 |

B. Effective connectivity parameters sig. predicted by gender alone

| | | | | | | | | |
|---|---|---|---|---|---|---|---|---|
| PCC to AI | Gender | .011 | | (1,575) = 6.615 | .010** | | | |
| | Gender | .021 | .010 | (4,572) = 3.088 | .016* | .128 | -.097 | .027* |
| | Dia BP | | | | | | -.057 | .324 |
| | Sys BP | | | | | | .014 | .815 |
| | BMI | | | | | | -.074 | .100 |
| mPFC to AI | Gender | .008 | | (1,579) = 4.578 | .033* | | | |
| | Gender | .013 | .005 | (2,578) = 3.714 | .025* | .093 | .084 | .043* |
| | BMI | | | | | | .070 | .093 |
| LIPC to PCC | Gender | .009 | | (1,576) = 5.143 | .024* | | | |
| | Gender | .015 | .006 | (3,574) = 2.800 | .036* | .180 | -.071 | .103 |
| | Dia BP | | | | | | -.023 | .696 |
| | Sys BP | | | | | | -.063 | .298 |
| RIPC to ACC | Gender | .008 | | (1,576) = 4.517 | .034* | | | |
| | Gender | .020 | .012 | (3,574) = 3.829 | .010** | .032* | -.054 | .215 |
| | Dia BP | | | | | | -.012 | .840 |
| | Sys BP | | | | | | -.106 | .079 |

C.1 Effective connectivity parameters not sig. predicted by the full model

| | | | | | | | | |
|---|---|---|---|---|---|---|---|---|
| PCC to mPFC | Gender | .000 | | (1,576) = .095 | .758 | | | |
| | Gender | .008 | .008 | (2,575) = 2.417 | .090 | .030* | -.016 | .710 |
| | Sys BP | | | | | | .095 | .030* |
| mPFC to RIPC | Gender | .000 | | (1,579) = .002 | .962 | | | |
| | Gender | .010 | .010 | (2,578) = 2.800 | .062 | .018* | -.005 | .910 |
| | BMI | | | | | | .098 | .018* |

| | | | | | | | | |
|---|---|---|---|---|---|---|---|---|
| mPFC to ACC | Gender | .001 | | (1,579) = .734 | .392 | | | |
| | Gender<br>BMI | .009 | .007 | (2,578) = 2.543 | .080 | .038* | .030<br>.087 | .474<br>.038* |
| RIPC to PPC | Gender | .001 | | (1,576) = .434 | .510 | | | |
| | Gender<br>Dia BP | .010 | .009 | (2,575) = 2.887 | .057 | .021* | -.012<br>-.097 | .771<br>.021* |
| DLPFC to AI | Gender | .000 | | (1,576) = .003 | .957 | | | |
| | Gender<br>Dia BP | .008 | .008 | (2,575) = 2.191 | .113 | .037* | -.012<br>.088 | .784<br>.037* |
| ACC to PPC | Gender | .000 | | (1,410) = .056 | .813 | | | |
| | Gender<br>BMI<br>HbA1c | .009 | .009 | (3,408) = 1.294 | .276 | .149 | -.014<br>.073<br>.054 | .783<br>.141<br>.280 |
| C.2 Effective connectivity parameters sig. predicted by the full model | | | | | | | | |
| AI to ACC | Gender | .000 | | (1,569) = .005 | .945 | | | |
| | Gender<br>Dia BP<br>Sys BP | .014 | .014 | (3,574) = 2.666 | .047* | .019* | .027<br>-.098<br>-.028 | .537<br>.092<br>.638 |
| AI to DLPFC | Gender | .001 | | (1,576) = .492 | .483 | | | |
| | Gender<br>Dia BP<br>Sys BP | .016 | .015 | (3,574) = 3.138 | .025* | .012* | .067<br>-.022<br>-.113 | .124<br>.705<br>.061 |
| AI to PPC | Gender | .006 | | (1,407) = 2.639 | .105 | | | |
| | Gender<br>Dia BP<br>Sys BP<br>BMI<br>HbA1c | .040 | .034 | (5,403) = 3.381 | .005** | .007** | .119<br>-.109<br>-.071<br>-.034<br>-.041 | .022<br>.114<br>.330<br>.527<br>.410 |

*Note*. **A.** The significant results, including the gender as predictor alone. **B.** Effective connectivity parameters significantly predicted by gender alone. As can be seen from Sig. F Change, the independent variables did not make a significant contribution to explaining the variance in the outcome variables, except for in RIPC to ACC, but for this regression none of the independent variables made a unique significant contribution. **C1.** Effective connectivity parameters where the full models were not significantly explaining the variance of the outcome variables. For all the regressions except for ACC to PPC the independent variables contributed to explaining more of the variance than gender alone. **C2.** Effective connectivity parameters where the full models were significantly explaining the variance of the outcome variables, and the independent variables explained significantly more than gender alone, but none of the independent variables made a unique significant contribution. *Abbreviations:* Dia BP = diastolic blood pressure; Sys BP = systolic blood pressure; BMI = Body Mass Index; HbA1c = glycated hemoglobin; PCC = posterior cingulate cortex; AI = anterior insula; mPFC = medial prefrontal cortex; LIPC/RIPC = left/right inferior parietal cortex; ACC = anterior cingulate cortex; DLPFC = dorsolateral prefrontal cortex; PPC = posterior parietal cortex; df reg = *degrees of freedom* regression; df res = *degrees of freedom* residual; beta = standardized coefficient

*$p < .05$, **$p < .01$.

*Supplementary Table 4.*

Non-significant Between BMI Group Differences for Effective Connectivity Parameters

| Connections | Sig. | $Chi^2$ |
|---|---|---|
| PCC | .398 | 1.846 |
| PCC to LIPC | .284 | 2.519 |
| PCC to AI | .314 | 2.319 |
| PCC to ACC | .127 | 4.125 |
| PCC to PPC | .691 | .744 |
| mPFC | .818 | .401 |
| mPFC to PCC | .461 | 1.550 |
| mPFC to LIPC | .199 | 3.225 |
| mPFC to AI | .446 | 1.616 |
| mPFC to ACC | .257 | 2.716 |
| mPFC to DLPFC | .157 | 3.697 |
| mPFC to PPC | .214 | 3.084 |
| LIPC to AI | .792 | .467 |
| LIPC to ACC | .993 | .014 |
| LIPC to DLPFC | .586 | 1.069 |
| LIPC to PPC | .328 | 2.231 |
| RIPC to PCC | .717 | .666 |
| RIPC mPFC | .824 | .386 |
| RIPC to LIPC | .944 | .115 |
| RIPC to AI | .507 | 1.358 |
| RIPC to ACC | .065 | 5.478 |
| RIPC to DLPFC | .648 | .868 |
| RIPC to PPC | .671 | .797 |
| AI | .709 | .688 |
| AI to PCC | .567 | 1.136 |
| AI to LIPC | .917 | .173 |
| AI to RIPC | .627 | .934 |
| AI to ACC | .861 | .300 |
| AI to DLPFC | .957 | .088 |
| AI to PPC | .096 | 4.683 |
| ACC | .368 | 2.001 |
| ACC to PCC | .474 | 1.494 |
| ACC to mPFC | .138 | 3.965 |
| ACC to LIPC | .871 | .275 |
| ACC to RIPC | .288 | 2.491 |
| ACC to AI | .446 | 1.614 |
| ACC to DLPFC | .775 | .509 |
| ACC to PPC | .196 | 3.255 |
| DLPFC | .648 | .867 |
| DLPFC to PCC | .233 | 2.916 |
| DLPFC to mPFC | .199 | 3.230 |
| DLPFC to LIPC | .080 | 5.043 |
| DLPFC to RIPC | .198 | 3.243 |
| DLPFC to AI | .570 | 1.125 |
| DLPFC to ACC | .127 | 4.132 |
| DLPFC to PPC | .694 | .730 |
| PPC | .124 | 4.174 |
| PPC to PCC | .092 | 4.779 |
| PPC to mPFC | .720 | .656 |
| PPC to LIPC | .315 | 2.308 |

| | | |
|---|---|---|
| PPC to RIPC | .834 | .363 |
| PPC to AI | .059[a] | 5.674 |
| PPC to ACC | .900 | .210 |
| PPC to DLPFC | .661 | .829 |

*Note.* Normal Weight (19-24 kg., N = 262), overweight (25-29 kg., N = 202) and obese (>30 kg., N = 117), Total N= 585, *degrees of freedom* = 2. [a] Close to significant group differences for PPC to AI. *Abbreviations:* PCC = posterior cingulate cortex; mPFC = medial prefrontal cortex; LIPC/ RIPC = left / right inferior parietal cortex; AI = anterior insula; ACC = anterior cingulate cortex; DLPFC = dorsolateral prefrontal cortex; PPC = posterior parietal cortex.

*$p < .05$

*Supplementary Figure 1.*

Frequency Distribution for BMI Groups

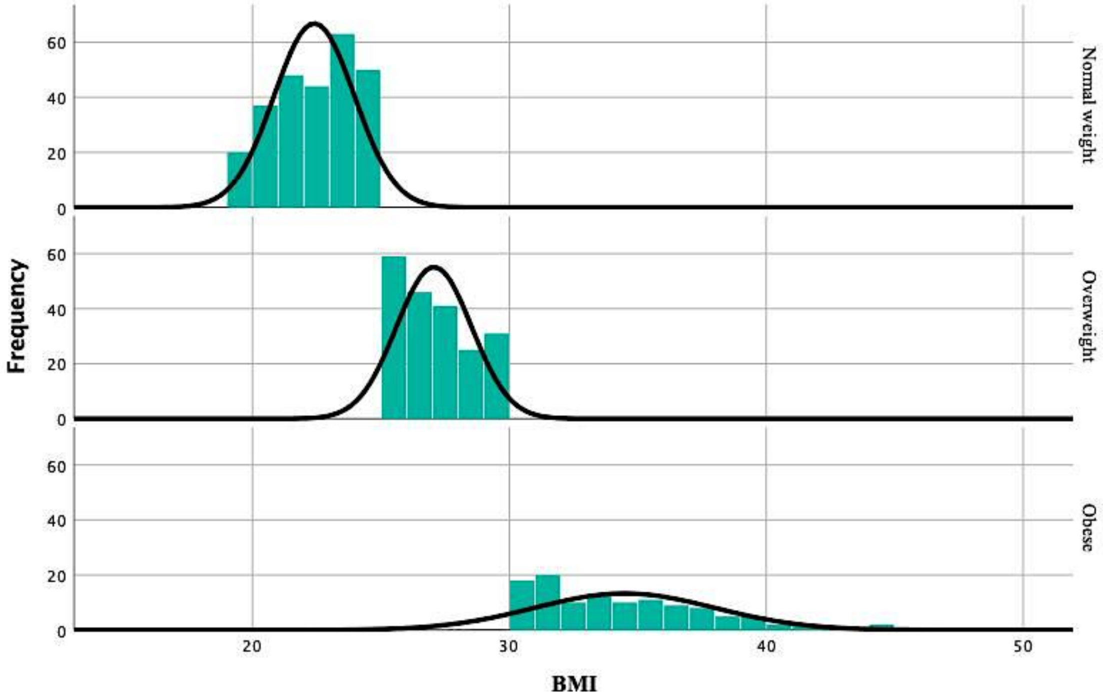


*Note.* Exhibits a non-normal distribution of BMI, and the shape of the distribution is not similar across the groups. Normal weight (19-24 kg., N=262), Overweight (25-29 kg., N=202) and Obese (>30 kg., N=117). The dependent variables did not show a similar distribution across groups and it was therefore considered most useful to report mean rank values for comparison between groups.
